# Structure-activity relationship of flavin analogs that target the FMN riboswitch

**DOI:** 10.1101/389148

**Authors:** Quentin Vicens, Estefanía Mondragón, Francis E. Reyes, Philip Coish, Paul Aristoff Judd Berman, Harpreet Kaur, Kevin W. Kells, Phil Wickens, Jeffery Wilson, Robert C. Gadwood, Heinrich J. Schostarez, Robert K. Suto, Kenneth F. Blount, Robert T. Batey

**Affiliations:** Department of Chemistry and Biochemistry, University of Colorado, 596 UCB, Boulder, CO 80309, USA; BioRelix Inc., 124 Washington St, Foxborough, MA 02035, USA; Aristoff Consulting LLC, 3726 Green Spring Drive, Fort Collins, CO 80528, USA; Dalton Pharma Services, 349 Wildcat Rd., Toronto, ON M3J 2S3, Canada; Kalexsyn, Inc., 4502 Campus Dr., Kalamazoo, MI 49008, USA; Xtal BioStructures, Inc., 12 Michigan Dr., Natick, MA 01760, USA

**Author notes:** Department of Biochemistry and Molecular Genetics, RNA BioScience Initiative, University of Colorado Denver School of Medicine, Aurora, CO 80045, USA. Thermo Fisher Scientific, 5350 NE Dawson Creek Drive, Hillsboro, OR 97124, USA. School of Forestry and Environmental Studies, Yale University, 195 Prospect Street, New Haven, CT 06511, USA. Rebiotix, Inc., 2660 Patton Road, Roseville, MN 55113, USA.

## Abstract

The flavin mononucleotide (FMN) riboswitch is an emerging target for the development of novel RNA-targeting antibiotics. We previously discovered an FMN derivative —5FDQD— that protects mice against diarrhea-causing *Clostridium difficile* bacteria. Here, we present the structure-based drug design strategy that led to the discovery of this fluoro-phenyl derivative with antibacterial properties. This approach involved the following stages: (1) structural analysis of all available free and bound FMN riboswitch structures; (2) design, synthesis and purification of derivatives; (3) *in vitro* testing for productive binding using two chemical probing methods; (4) *in vitro* transcription termination assays; (5) resolution of the crystal structures of the FMN riboswitch in complex with the most mature candidates. In the process, we delineated principles for productive binding to this riboswitch, thereby demonstrating the effectiveness of a coordinated structure-guided approach to designing drugs against RNA.

**GRAPHICAL ABSTRACT:** Exploring the chemical structure landscape of FMN riboswitch binders.

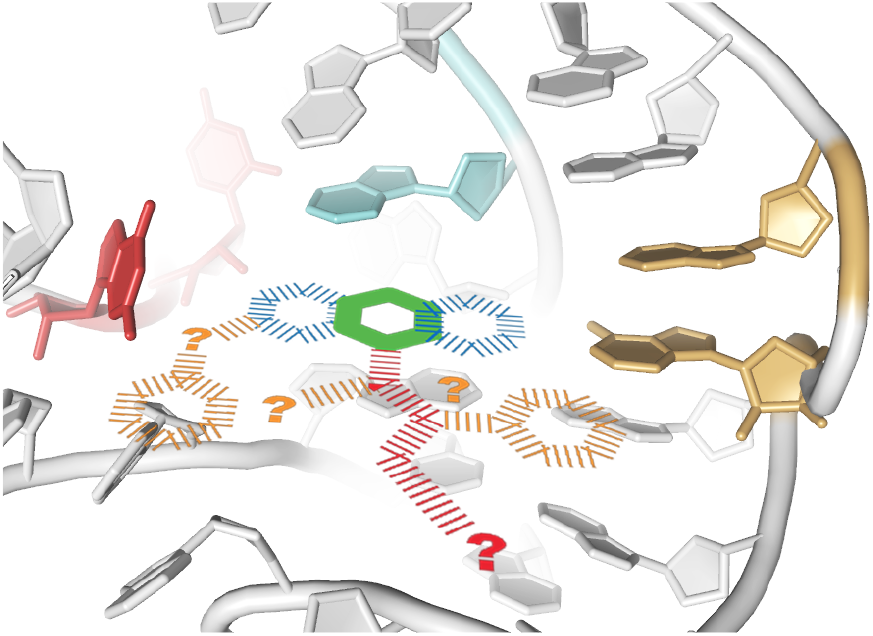

## INTRODUCTION

Since seminal work on antibiotic-RNA complexes in the 1980s-90s, RNA has been recognized as a promising therapeutic target for small molecules.^1–2^ At least half of the known families of antibiotics target ribosomal RNA, including linezolid, one of the most recently discovered antibiotics.^3^ Since the year 2000, three-dimensional structures of ribosome-antibiotic complexes solved using X-ray crystallography and cryo-electron microscopy have helped us decipher drug-RNA binding principles.^4^ Today, many companies —including pharmaceutical giants like Merck, Pfizer, and Novartis— are running programs aimed at targeting RNA with small molecules.^5–6^

Biosensors mostly found in bacteria and called “riboswitches” were recognized soon after their discovery as promising RNA drug targets, mostly because they consist of structured RNA elements that regulate the expression of genes essential for survival/virulence of some medically important pathogens through the binding of small molecules or ions.^7–9^ Disrupting molecular switches is a proven strategy for achieving inhibitory bioactivity,^10^ including documented examples of riboswitches that can productively bind natural products and anti-metabolites. For example, sinefungin binds to the S-adenosyl methionine (SAM) riboswitch,^11^ antifolates can regulate the tetrahydrofolate riboswitch,^12^ and the *thi*-box riboswitch was crystallized in the presence of a variety of thiamine pyrophosphate analogs.^13^ However, analog binding is not synonymous with antimicrobial action *in vivo*. For instance, although lysine analogs bind to the lysine riboswitch,^14^ their antibacterial activity actually mostly stems from their incorporation into nascent proteins.^15^ Notably, a case where binding of a compound has an antimicrobial effect is illustrated by roseoflavin (RoF) —produced by *Streptomyces davawensis*—^16^, whose antibacterial action comes from binding of its phosphorylated form (RoFMN) to the flavin mononucleotide (FMN) riboswitch.^17^ RoFMN binding downregulates the expression of downstream genes,^17^ which offers a compelling evidence that riboswitches are generally valid targets for drug design efforts.^18^

For structure-guided design of novel riboswitch-targeting compounds, the three-dimensional architecture of the complex between the riboswitch and its natural cognate ligand is the starting point for exploring chemical space through optimizing the pharmaceutical properties of the ligand.^19–21^ For example, this strategy was used to design purine analogs that had antimicrobial action by binding to the purine riboswitch.^19^ Additional structure-function characterizations guided the design of modified pyrimidines that could also be accommodated in the binding pocket of the purine riboswitch, in order to interfere with gene expression.^20, 22^ These successful efforts illustrate that reaching ligand-specific pharmacochemical goals leads to superior therapeutic compounds.

A key aspect of structure-guided design approaches is that they do not solely rely on visualizing ligand-bound states, but also on interrogating the ensemble of unbound states, using techniques such as in-line probing,^8, 23^ Selective 2’ Hydroxyl Acylation analyzed by Primer Extension (SHAPE),^24^ small angle X-ray scattering,^25^ fluorescence,^26^ NMR,^27^ and most recently X-ray free electron laser.^28^ Precisely mapping local conformational changes associated with ligand binding is in fact critical for identifying ligand positions that can be modified, while maintaining activity. The bioactivity of the resulting compounds is in turn tested using the same techniques. In the case of the purine, SAM, and FMN riboswitches, these methods suggested that ligand recognition occurs by conformational selection of a preformed structure, rather than through a more extensive allosteric mechanism.^24–25, 29–30^ Structure-guided drug design is particularly promising in such scenarios, because the architecture of the active site as seen in the bound structure is similar to that encountered by a ligand upon binding.

Herein, we present the main pathways of a medicinal chemistry strategy for optimizing RoFMN that led to the discovery of 5FDQD (5-(3-(4-fluorophenyl)butyl)-7,8-dimethylpyrido[3,4-b]-quinoxaline-1,3(2H,5H)-dione; Figure 1A), a synthetic fluoro-phenyl analog of RoFMN that binds to the FMN riboswitch of *Clostridium difficile*. Importantly, 5FDQD was shown to be bactericidal toward *C. difficile* without perturbing the host mouse microbiome.^31^ Furthermore, the frequency of developing resistance against 5FDQD is low (< 1 x 10^-9^) ^31^. Using a combination of chemical probing techniques and transcription termination assays, we characterized the contribution to RNA binding and regulatory activity of various FMN and RoFMN synthetic analogs. The structures of three of the most promising compounds were determined using X-ray crystallography. In addition, we performed a meta-analysis of known ligands that target the FMN riboswitch,^30, 32–33^ including Merck’s ribocil, an unnatural ligand with a novel chemical scaffold^32^. Principles for designing effective drugs could be derived, so that bioavailability, binding to this riboswitch, and efficiency are not compromised. Overall, this work further establishes the FMN riboswitch as a powerful model system for understanding how to target RNA.

**Figure 1.**
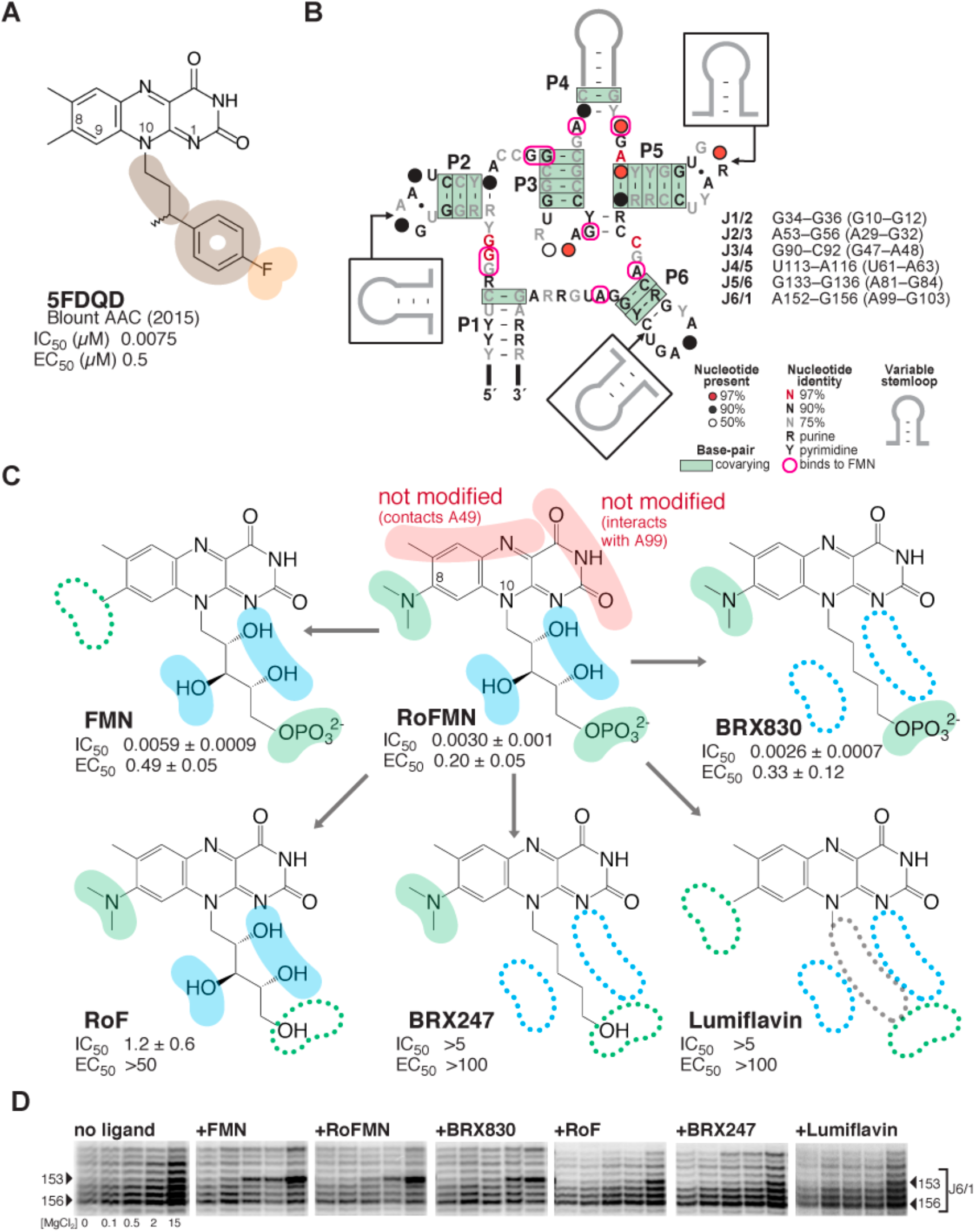
Roseoflavin mononucleotide at the center of a medicinal chemistry optimization strategy that led to the discovery of synthetic analogs with potent activity and selectivity. (A) Chemical structure of 5FDQD (5-(3-(4-fluorophenyl)butyl)-7,8-dimethylpyrido[3,4-b]-quinoxaline-1,3(2H,5H)-dione), a potent inhibitor of the FMN riboswitch. Color-coding for functional groups introduced during SAR study: orange, negatively charged/polar group; tan, hydrophobic group. IC_50_, half maximal inhibitory concentration as measured by in-line probing; EC_50_, half maximal effective concentration in transcription termination assays.^31^ Note that all values for IC_50_ and EC_50_ in subsequent figures are given in units of μM. (B) Secondary structure of the FMN riboswitch showing sequence and structure conservation among bacteria. Residues that interact with flavin-bearing ligands in crystal structures are circled in pink.^30, 33^ The list of the six joining regions is indicated, with numbering for *B. subtilis* and *F. nucleatum* (in parenthesis). (C) Dissecting functional positions 8 and 10 of roseoflavin mononucleotide (RoFMN) over the course of a structure-activity relationship (SAR) study of the FMN riboswitch. Color-coding for functional groups: red, left unaltered during SAR study; green, primary focus of SAR study; blue, secondary focus. IC_50_ and EC_50_ calculated as for 5FDQD (see Methods; Tables S1 and S2; Figure S1). (D) Comparative banding pattern of SHAPE chemical probing within the J6/1 joining region, which serves as an indicator for ligand binding.^30^ [MgCl_2_] tested were: 0, 0.1, 0.5, 2.0 and 15.0 mM. Arrowheads: residues of interest within J6/1. The gels were aligned in SAFA^55–56^ (full unaltered SHAPE gels shown in Figure S2).

## RESULTS AND DISCUSSION

### Design rationale

Our rationale for optimizing RoFMN stemmed from the following challenges. First, until after this project was completed,^34–35^ roseoflavin was thought to enter bacteria only via an active riboflavin transporter specific to Gram-positive bacteria.^36–37^ This could limit intracellular concentrations of roseoflavin, thereby restricting its potency and activity spectrum. In addition, since these riboflavin transporters are not essential, their mutation could render bacteria resistant to roseoflavin. Second, roseoflavin requires intracellular phosphorylation to exert antibacterial activity as RoFMN,^17^ as suggested by in-line probing and fluorescence-based assays that demonstrated a ~1,000-fold decrease in binding affinity when the phosphate group is removed.^23, 33^ The requirement for phosphorylation also constitutes another avenue for resistance to emerge. Moreover, RoFMN antibacterial activity could be self-limiting, if growth inhibition decreases the effectiveness of roseoflavin phosphorylation. Third, the need for roseoflavin to be recognized and activated by multiple proteins required for its transport and phosphorylation imposes additional structural and functional constraints on the ligand. Finally, roseoflavin is rapidly cleared from plasma (K.F.B., unpublished results). Given these challenges, our pharmacochemical goals were to identify RoFMN analogs that could be passively transported into bacteria and not require phosphorylation to be active.

The first optimization stage was to examine the interaction of FMN and roseoflavin with FMN riboswitches for key recognition motifs and behaviors that new analogs would need to maintain or replace.^30, 33^ Four elements seemed important: [1] a pseudo-base pair involving A99 and the uracil-like face of the isoalloxazine ring system; [2] a metal ion-mediated interaction involving the FMN phosphate and four of the six joining regions; [3] aromatic stacking between A48 (within joining (J) regions J3/4), the isoalloxazine moiety, and A85 (adjacent to J5/6); and [4] four of the six joining regions interacting with the ligand in crystal structures^30, 33^ (Figure 1B) undergo significant conformational change upon binding, two of which maintain a structure of the binding pocket that is conducive to ligand binding: J4/5 (which through G62 is part of a metal-mediated interaction involving the phosphate group at position 10 of FMN), and J3/4 (which through A48 is in the vicinity of U61 and position C8 of FMN, a natural site for derivatization (Figure 1C)^38^).^30^ Furthermore, the hydroxyl groups of the ribityl-phosphate chain at position 10 of the isoalloxazine ring system do not contact RNA and their contribution to binding is minimal.^23, 33^ Hence, although the ribityl group forms a hydrogen bond to O6 of G11, it appears to merely direct a negatively charged group to a critical location. Its hydroxyl groups could likely be removed without altering activity, while potentially significantly improving passive membrane permeation. From these observations, it was concluded that positions 8 and 10 on the isoalloxazine ring system would be the most promising in the search for synthetic flavin analogs that bypass phosphorylation requirements.

### Step-by-step breakdown of the ribityl group off position 10 on the flavin heterocycle

To assess whether hydroxyl groups on the ribityl chain at position N10 of the isoalloxazine heterocycle were dispensable, we carried out a systematic dissection of the contributions of each chemical element at that position. To that end, compounds were purchased or synthesized (see SI Methods) that progressively stripped functional groups off the N10 chain (Figure 1C). BRX247 and BRX830 were the endpoints of this series, in which all hydroxyl groups on the ribose were removed to yield an alkyl chain. In BRX247, the terminal phosphate group was removed.

We comparatively assessed the effect of these compounds on the structure and the activity of the *Bacillus subtilis ribD* FMN riboswitch (Figure S1A) bound to these derivatives using inline probing,^23, 39^ SHAPE, ^30, 40^ and transcription termination assays.^31^ FMN, roseoflavin, and lumiflavin were used as controls.^17, 23, 33^ These data largely recapitulated earlier findings on the relative importance of the ribityl and phosphate groups, in support of RoFMN being the active form of roseoflavin in cells.^17^ In addition, we showed that an alkyl chain could substitute for the ribityl chain (compare FMN and BRX830; Figure 1C). Further, SHAPE analysis revealed that BRX830, but not BRX247, promoted structural changes in the binding pocket similar to that of RoFMN, suggestive of productive binding to the riboswitch (Figure 1D). This observation confirmed the need for a negatively charged group to organize the binding core of the RNA, while also encouraging the design of alternatives to the ribityl chain and potentially the phosphate group at its end.

### Synthetic ligands with superior potency to roseoflavin mononucleotide

With the next series of compounds, we explored whether the phosphate group could be functionally replaced by a less labile negatively charged group. When the phosphate group of BRX830 was replaced by a carboxylic group (leading to the heptanoic acid side chain at position 10 in BRX857; Figure 2A), a 100-fold decrease in IC_50_, and a 6-fold loss in EC_50_ were observed. Shortening the alkyl chain as in BRX897 (from 6 to 4 carbon atoms, leading to a pentanoic acid side chain; Figure 2A) resulted in slightly lower binding affinity, and a further 10-fold reduction in activity. However, the riboswitch was tolerant of various chain lengths within a 5–7 carbon-atom range (affinities varied between 0.2-0.9 μM; EC_50_ for 4–7 carbon atoms were ranked as follows: C6 > C5 ≫ C4,C7; data not shown). Both BRX857 and BRX897 partially induced a roseoflavin-like conformational changes at the binding site, as indicated by SHAPE (Figure 2B). Notably, removing the *N*-dimethyl group at position 8 led to a 10-fold improvement in affinity and a four-fold improvement in activity (compare BRX1027 to BRX857; Figure 2A). Other replacements of the phosphate group were less effective for binding than carboxylic acid (e.g., IC_50_ (nitrile) ~ 4.5 μM; data not shown). Interestingly, moving the heptanoic acid chain from position N10 to positions N1 or C9 on the isoalloxazine heterocycle led to compounds that showed improved permeability as tested using a parallel artificial membrane permeability assay,^41^ but weaker functional activity (for BRX1027, permeability index (pe) = 0.87; for heptanoic acid at N1: IC_50_ = 0.7 μM, EC_50_ = 8 μM, pe = 8.4; for heptanoic acid at C9: IC_50_ = 0.33 μM, EC_50_ = 1.7 μM, pe = 5.4; data not shown). In summary, BRX1027 displayed physicochemical properties superior to those of the natural ligands, but its effectiveness in promoting transcription termination was weak. BRX1027 thus provided a novel structural framework, where potential functionalization sites could be incorporated both at positions 8 and within the heptanoic acid chain.

**Figure 2.**
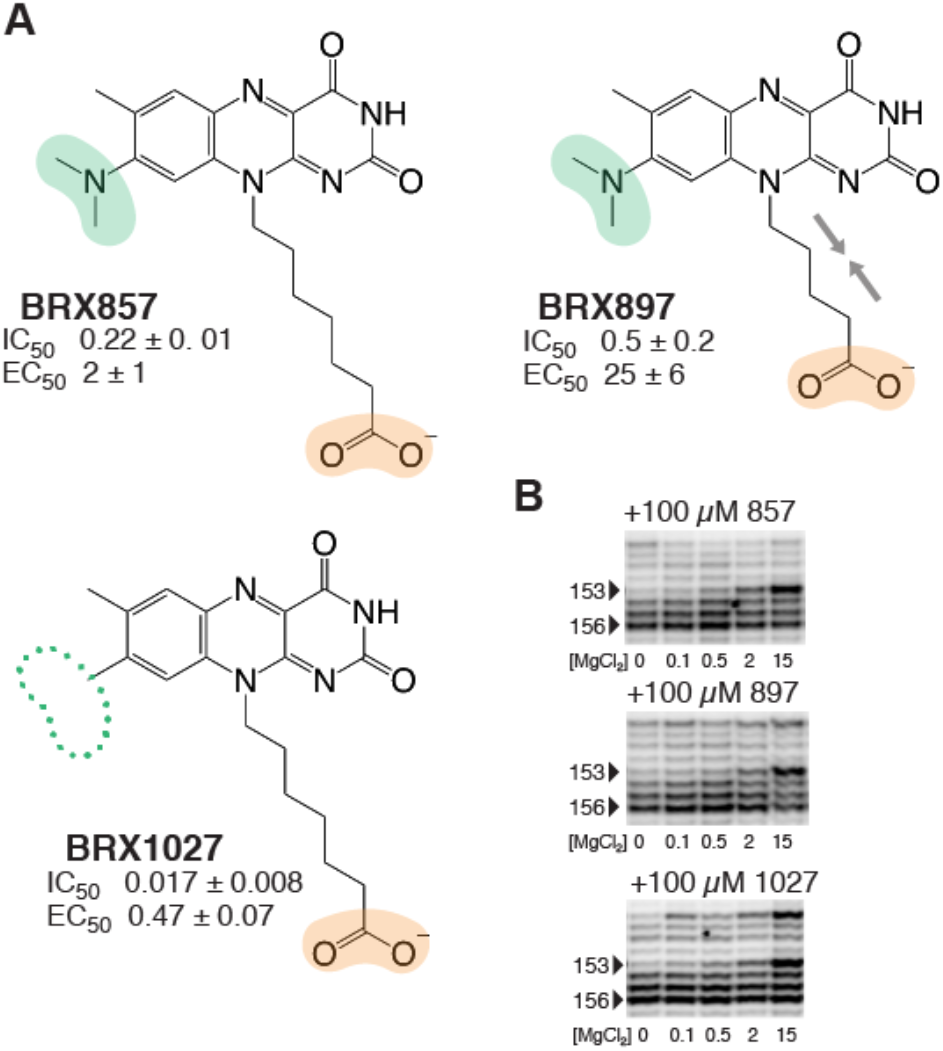
Drug leads in which the phosphate group was replaced by a carboxylic acid. (A) A carboxylic group in place of the phosphate group only results in a minor drop in binding and activity. Color-coding for functional groups: green, as in Figure 1; orange, negatively charged/polar group introduced during SAR study (raw and normalized data in Tables S1 and S2; Figure S1). (B) Banding pattern of SHAPE chemical probing within the J6/1 joining region for compounds shown in panel (A), as in Figure 1 (full unaltered SHAPE gels shown in Figure S2).

The next aspect of this SAR study examined the effect of adding hydrophobic and heterocyclic groups at positions 8 and 10 of the isoalloxazine ring system in BRX1027. Such ring systems make for trademarks of successful designer drugs.^1, 42–43^ For example, adding a cyclopentyl ring at the N8 position of BRX857 (as in BRX1151) partially restored high affinity binding while improving transcription termination by a factor of 10 (Figure 3A). Also, although the introduction of an amine within the heptanoic acid chain leads to a decrease in affinity and activity (BRX931; Figure 3A), it opened the possibility of introducing various functionalizations, such as a piperidine (as in BRX1224; Figure 3A). Going from BRX931 to BRX1224, binding affinity had improved by 26-fold and activity by 225-fold (Figure 3A). These effects were not necessarily additive though, as demonstrated by BRX1072, which had a lower affinity than BRX1151 and BRX1224, despite possessing both a piperidine as in BRX1224 and a cyclopropyl at position 8 (Figure 3A). BRX1151 but also to some extent BRX1072 and BRX1224 were successful at promoting FMN-like conformational changes of the binding site (Figure 3B), which encouraged us to solve the structure of BRX1151 bound to the riboswitch (see below).

**Figure 3.**
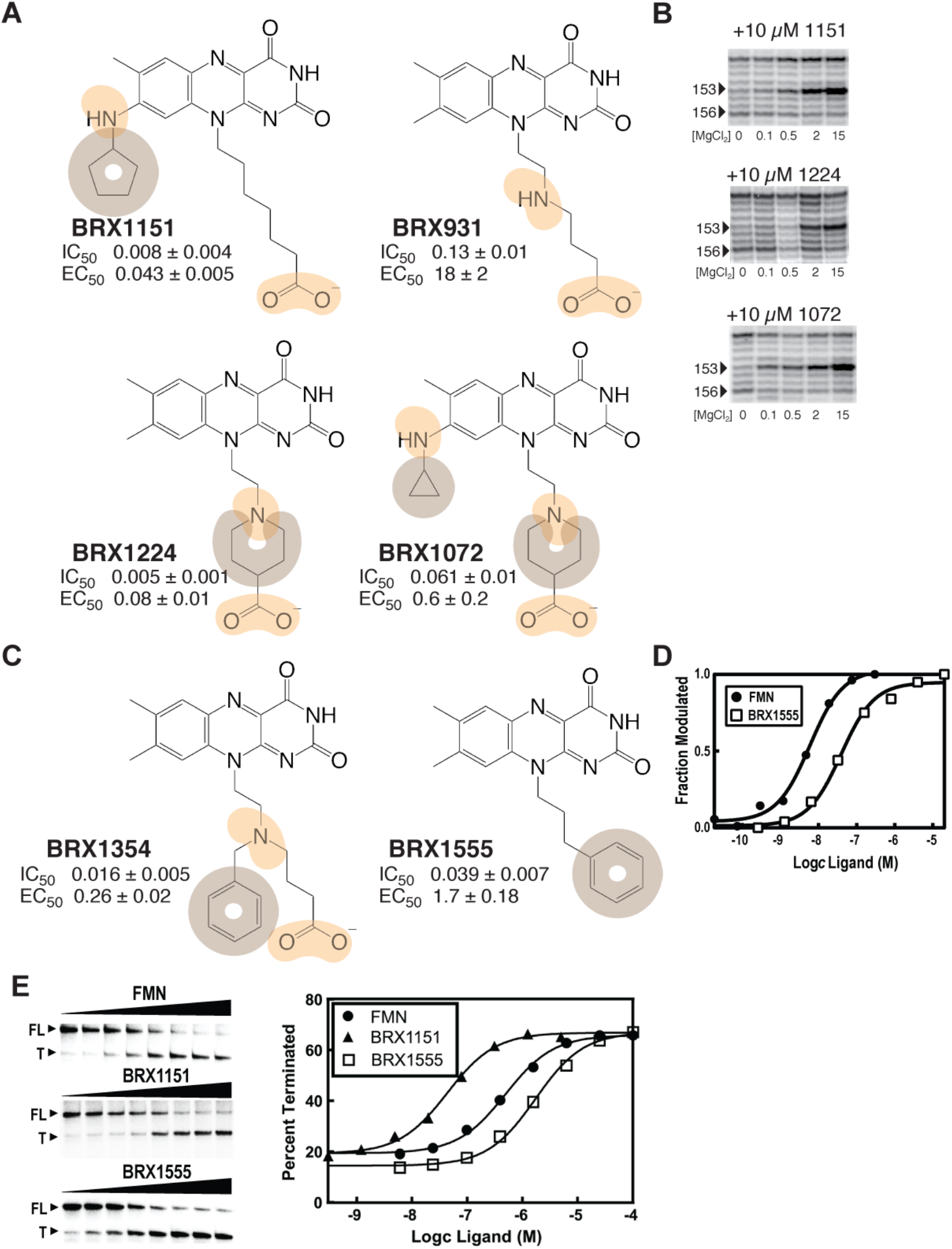
Developing mature drug leads by adding hydrophobic groups and heteroatoms. (A) Hydrophobic groups and heterocycles at positions 8 and/or 10 restore partial binding and efficacy. Color-coding for functional groups: orange, same as above; tan, hydrophobic. (B) SHAPE banding pattern for BRX1151 and BRX1072. (C) Ligands with aromatic side chains off position 8. (D) Plot of the average of the normalized fraction of RNA cleaved at G136 versus FMN/BRX1555 concentration. Curves indicate the best fit of the data to an equation for a two-state binding model (raw data shown in Figure S1). (E) Polyacrylamide gel analysis of a transcription termination assay using BRX1151, BRX1555 and FMN (left). Plot depicting the fraction of transcripts in the terminated form versus ligand concentration (right). Curves indicate the best fit of the data to a two-state model for ligand-dependent termination activity (raw and normalized data in Tables S1 and S2; Figure S1; full unaltered SHAPE gel shown in Figure S2).

At that point, BRX1151 was the most attractive candidate to emerge from our study, with IC_50_ and EC50 values that were sufficiently low to warrant bioactivity. Yet, although less labile than a phosphate group *in vivo*, a negatively charged carboxylic group still depends on a transporter to cross bacterial membranes, representing a major limitation to bioavailability. Thus, BRX1151 merely served as a proof of principle that the phosphate group responsible for the activity of (Ro)FMN and BRX830 could be replaced by another negatively charged group. To generate a derivative that would be effective as a drug, we further explored the derivatization of BRX1151 to bypass the need for a negatively charged functional group.

### Aromatic hydrocarbons to the rescue: the discovery of BRX1555

To circumvent critical issues of having a derivative with a carboxylic group, an approach consisting in introducing hydrophobic groups proved most rewarding. Adding a phenyl moiety to BRX931 through an amine inserted in the alkyl chain led to a recovery in binding affinity of the same order of magnitude as that seen in the compounds with cyclopentyl or piperidine, i.e. only about 2-3 times weaker than FMN (BRX1354; Figure 3C). This addition also had the effect of restoring transcription termination inhibition to the level of FMN. Bypassing the carboxylic group but maintaining an aromatic group at the end of the alkyl chain resulted in BRX1555, a compound with a binding affinity of 39 nM, and an EC_50_ of 1.7 μM (similar to that of BRX857; Figures 2 and 3C, D). Increasing the length of the alkyl chain in BRX1555 by one carbon atom decreased transcription termination efficiency by a factor of 10 (data not shown). Other alternatives to the carboxylic group combining heteroatoms and aromatic rings were less effective (e.g., EC_50_ (-pentyl thiazolidinedione) = 5.1 μM; EC_50_ (-piperidinyl acyl sulfonamide) = 23 μM; data not shown). Worthy of note, tert-butyl and phenethyl carboxylate derivatives that could be used as prodrugs consistently displayed weak activity in microbiology assays (e.g., MIC ≤ 16 μg/mL against *Staphylococcus aureus* NRS384), but were unstable (plasma levels of 7.6 μg/mL (tertbutyl), 0.08 μg/mL (phenylethyl), and under quantifiable limits (tertbutyl with C8-chloro substitution), as measured 5 min after administering a 5 mg/kg IV dose to rodents; data not shown).

Fortunately, the promising IC_50_ and EC_50_ values of BRX1555 were accompanied by a weak but consistent activity against several strains of *Staphylococcus aureus* (laboratory strain RN4220; Rosenbach, ATCC #29213; multi-resistant NRS384) and *Streptococcus pneumoniae* (Chester, ATCC #49619; Table S3; data not shown). Further modifications of BRX1555 at position 8 (e.g., removal of the methyl group, chloro substitution, addition of propylene glycol to N8) led to compounds with improved solubility, cytotoxicity, but not potency (Table S3; data not shown). Subsequent efforts exploring the BRX1555 scaffold led to the identification of 5FDQD —originally called BRX2102— which retained the BRX1555 skeleton with the addition of a methyl group on the alkyl chain and a fluorine substituent on the phenyl moiety.

### Substituents at positions 8 and 10 confer selectivity for certain bacterial strains

While exploring the effect on binding of adding substituents at position 8, we noticed an effect on selectivity. For example, BRX1027 was compared to BRX1070 (similar to BRX1027 but possessing a cyclopropyl substituent at position N8; Figure 4A) for binding to the FMN riboswitch from four bacterial strains, including *Staphylococcus aureus* and *Staphylococcus epidermidis,* using in-line probing assays. Both *S. aureus ypaA* and *S. epidermidis* were sensitive to a different extent to RoFMN (MIC_50_ (*S. aureus*) = 8 μg/mL; MIC_50_ *(S. epidermidis)* = 32 μg/mL; data not shown). The results of this comparative analysis were that although FMN and BRX1070 would bind similarly to all tested riboswitches, BRX1027 would not (Figure 4B). The affinity of BRX1027 for the *S. aureus* riboswitches was ~10-fold lower than that to *B. subtilis*, and ~20-fold lower to *S. epidermidis* (Figure 4B). A 35-fold decrease in affinity was also noticed for a halogenated derivative of BRX1555 (i.e., BRK1675; Figure 4A) between *B. subtilis* and a *C. difficile-specific* FMN riboswitch (i.e., CD3299; Figure 4B).^44^ Similarly, SHAPE mapping revealed poor binding of BRX1027 to the *Fusobacterium nucleatum* FMN riboswitch, in comparison to FMN, BRX857 and BRX1151 (Figure 4C). Together, these observations suggested that to target a specific strain, a substituent would need to be present at position 8 when a carboxylic acid is at position 10. Interestingly, in the context of BRX1555 (Table S3), BRX1675 (Figure 4) and 5FDQD,^31^ selectivity was achieved in absence of a substituent at position 8, emphasizing the interplay between the nature of a substituent at position 8, and that at position 10.

**Figure 4.**
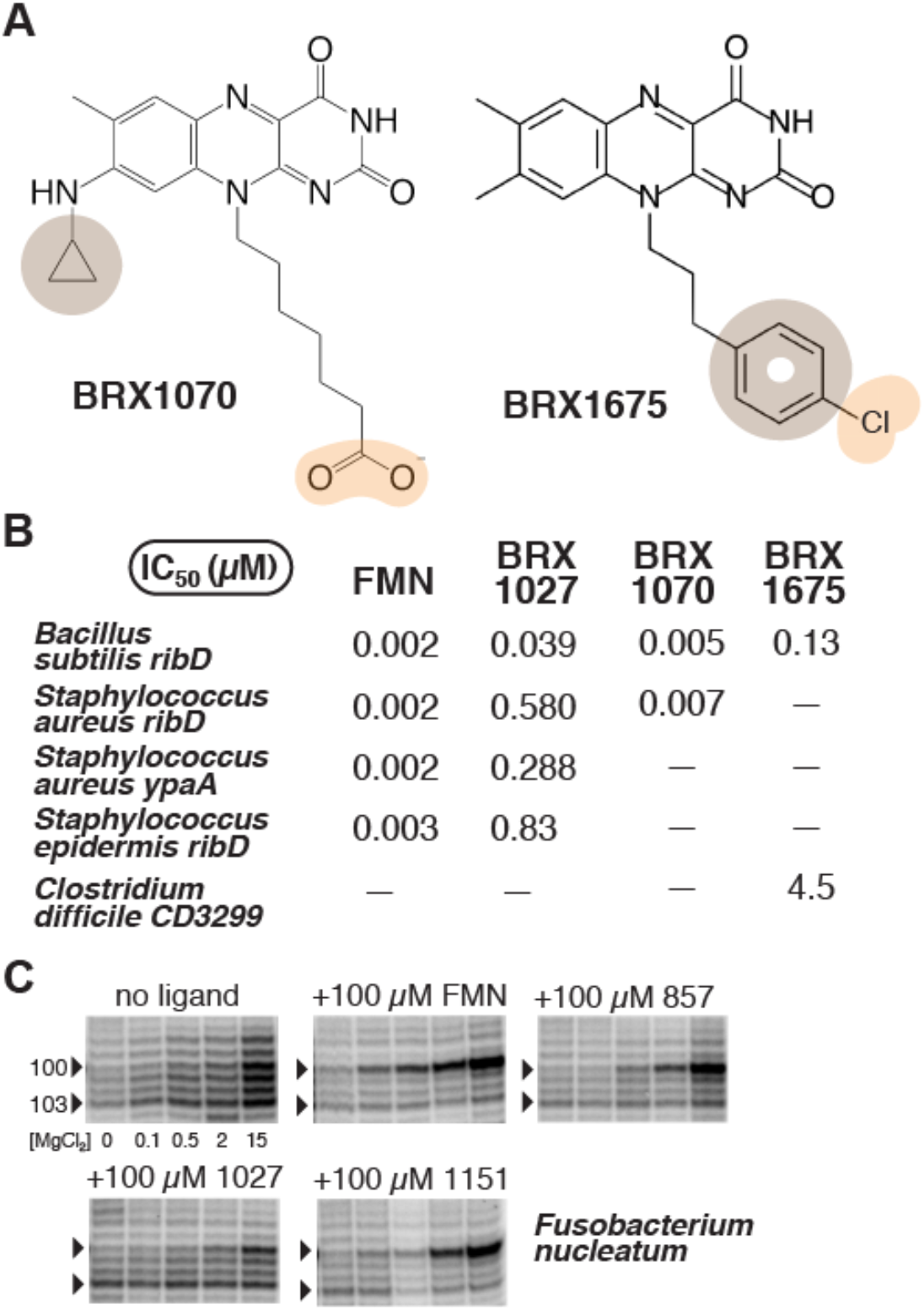
Selectivity of flavin analogs. (A) Chemical structures of BRX1070 and BRX1675. (B) IC_50_ values (μM) from in-line probing assays using FMN, three BRX compounds, and five different riboswitch sequences. (C) Mg^2+^-dependent probing of the *F. nucleatum* riboswitch with a series of BRX compounds. Residues 100 and 103 within the J6/1 joining region are marked by arrows (full unaltered SHAPE gel shown in Figure S2).

In order to gain selectivity for a particular flavin-based drug, the sequence of J4/5 (residues 61–63 in *F. nucleatum*) may need to be taken into account, as J4/5 differentially interacts with position 8 and position 10 substituents depending on the ligand (see next section). While the sequence of J4/5 is UGGA in *B. subtilis*, it is UGA in *F. nucleatum*, CUGA in the two *Staphylococcus* strains, and UUGA in *C. difficile*. Although this interplay in species selectivity had not been reported, it is compatible with earlier findings.^17^ For example, binding characteristics are similar whether roseoflavin binds to the FMN riboswitch from *B. subtilis* or from *S. davawensis* (J4/5 is UUGA), although it was proposed that the phosphorylated form of roseoflavin discriminates between the two species in a manner different from FMN.^17^ It is the combination of such observations about affinity, *in vitro* and *in vivo* activity, as well as selectivity, that eventually led to the design of 5FDQD, which is selectively active against *C. difficile* in the mouse gut.

### Distinct ligand-RNA interactions support a conserved architecture of the binding site

To gain insights into how functional groups added at positions 8 and 10 were accommodated in the binding site of the FMN riboswitch, we solved the crystal structures of BRX1151, BRX1354 and BRX1555 bound to the aptamer domain of the FMN riboswitch from *F. nucleatum*, using previously published strategies.^30, 33^ The overall RNA folds were similar for the three ligands and they supported a similar architecture of the binding site (Figure S3A). In all complexes, the isoalloxazine ring system intercalates between A48 and the G98-A85 pair, and the uridine-like moiety forms a pseudo-Watson-Crick base pair with A99, as in the FMN complex (Figure 5A–D; Figure S3B–D). Local structural differences occurred for the backbone atoms at J4/5, as previously reported for various ligands bound to the riboswitch (Figure S3A).^30, 33^ In short, crystal structures confirmed the structural similarity that was suggested by in-line probing and SHAPE data.

**Figure 5.**
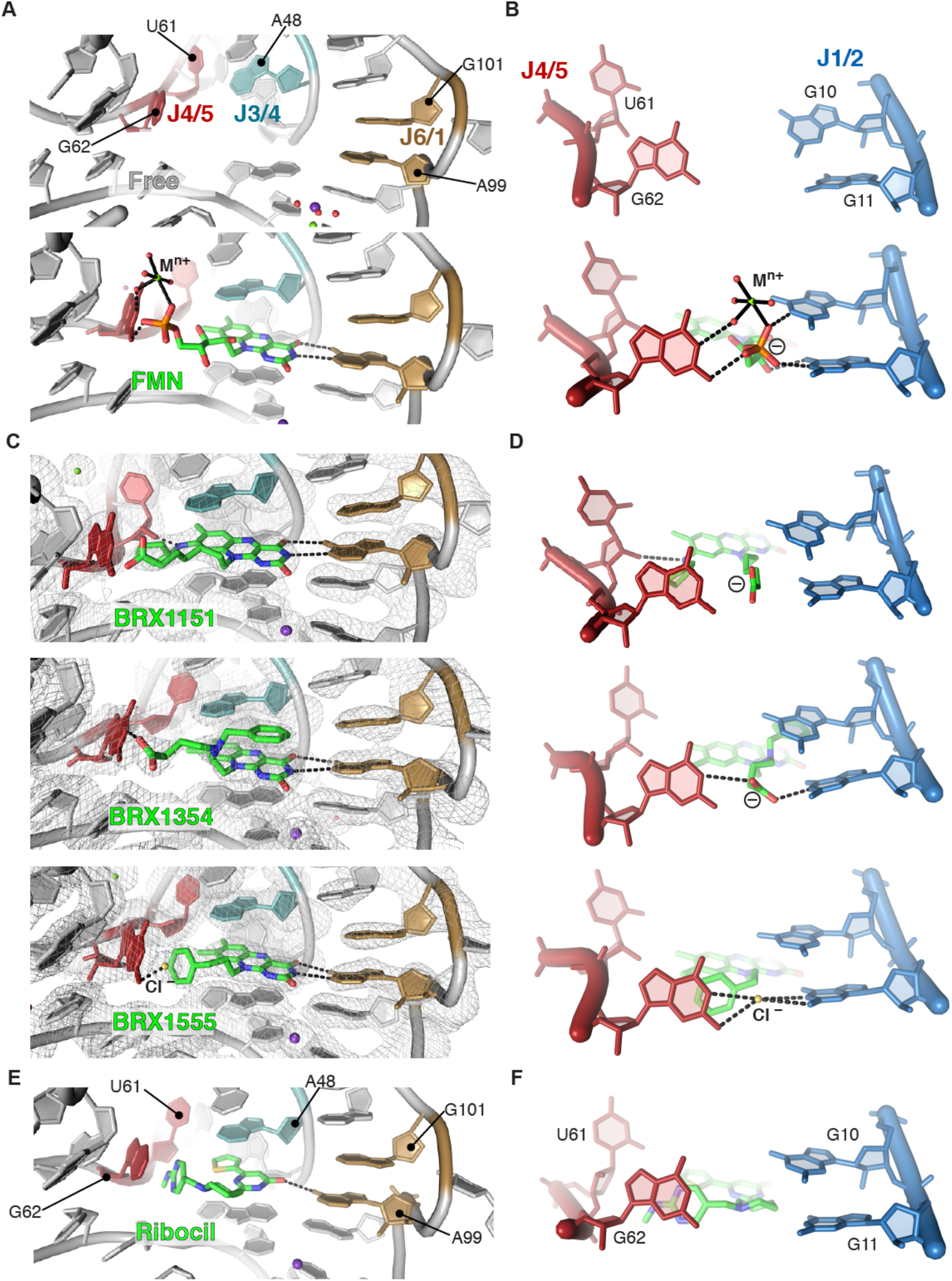
Visualizing how different chemical scaffolds promote binding to the FMN riboswitch. (A) Organization of the binding site in absence of ligand (PDB ID 2YIF) and in presence of FMN (PDB ID 2YIE).^30^ (B) Close-up on J1/2 and J4/5. (C) 2F_o_-F_c_ maps of three roseoflavin analogs (map contour: 1.0 σ), showing intercalation of the isoalloxazine moiety and its pseudo-Watson-Crick pairing with A99. J1/2 and J2/3 are omitted for clarity. (D) Plasticity of interaction types in the region involving J1/2 and J4/5. (E) Same view as in (A, C) for the FMN-riboswitch bound to ribocil (PDB ID 5C45).^32^ (F) Same view as in (B, D) for the ribocil-bound riboswitch.

An unexpected aspect of these structures is how the various modifications are accommodated in the binding pocket. The carboxylic group at the end of the alkyl chain (in BRX1151 and BRX1354) interacts with G62 in a similar manner to the phosphate group at the end of the ribose chain in FMN (Figure 5C, D). G62 had been identified as one of the four guanosines that interact with the phosphate group of FMN and a magnesium ion in the co-crystal structure,^33^ as well as belonging to one of the two joining regions (J3/4 and J4/5) that undergo conformational change upon binding.^30^ No metal ion was detected though in the case of the BRX complexes, suggesting a magnesium ion is dispensable for binding efficiency, at least in the absence of a phosphate group. In BRX1555, which does not contain any negative charge, the phenyl moiety at the end of the alkyl chain is stacked against G62 (Figure 5C, D). Within that structure, a strong density peak between G62 and G11 was best modeled as a chloride ion, consistent with the presence of a negative charge in that area in the other complexes (Figure S5E). However, further experiments would be needed to confirm the nature of the putative chloride ion. Also, the presence of the substituted —and likely protonated— amino group at position 8 of the isoalloxazine system in BRX1151 results in a hydrogen bond to the 2’-OH of U61 from J4/5 (Figure 5C). Finally, the phenyl moieties that were added to BRX1354 and BRX1555 adopt different orientations: in BRX1354 the ring stacks against the isoalloxazine moiety while in BRX1555 it stacks against G62 (Figure 5C, D). This variability in tolerating a similar functional group is not what we would have predicted from the chemical structures.

Overall, this select structural analysis of some of the most mature BRX compounds illustrated how the FMN riboswitch can adjust to recognizing ligands with a variety of functional groups, consistent with the flexibility reported by SHAPE for U61/G62 (J4/5) and G10/G11 (J1/2) in the free state ensemble.^30^ What these new structures further illustrate is that in addition to stacking against A48 (J3/4) and interacting with A99 (J6/1), some form of interaction with G62 is critical for productive binding (Figure 5A–D). Interestingly, the structure of ribocil, a compound identified in high-throughput screening supports that conclusion as well.^32^ Although its chemical structure is distinct from FMN relatives, its pattern of interactions within the binding pocket follows similar rules (Figure 5E, F). In particular, the 2-methylaminopyrimidine moiety of ribocil stacks against G62 in the crystal structure, in a manner that is consistent with what we observe for the phenyl ring of BRX1555 (Figure 5C, F). Further highlighting the critical role of the interaction between G62 and a ligand, it was proposed that disruption of this interaction was responsible for loss of FMN and ribocil binding in a riboswitch mutant in which G62 and adjacent residues had been deleted.^45^

### Making a case for integrated medicinal chemistry strategies

The discovery of 5FDQD exemplifies how an effective and selective drug can be designed by leveraging biochemical, functional and structural data about an RNA target.^31^ In this process, a key is to reach a confluence between the positions of a drug lead that appear modifiable according to available biochemical and structural data, and the elements that seem to limit effectiveness in the eyes of medicinal chemists. The FMN riboswitch is currently one of the few RNAs—together for example with the ribosomal decoding site^46–49^—for which natural antibacterials (e.g. roseoflavin),^17^ synthetic compounds (5FDQD and this work) and entirely new scaffolds (ribocil^32^, WG-3^50^) have been extensively studied, making it a particularly attractive system for improving our ability to target RNA. At least, the comparative analysis of the various ligands that bind to this riboswitch shows that exploring a vast chemical space outside of the natural variations seen between cognate ligands is not a vain endeavor.

The SAR approach in this study revealed several key aspects of designing therapeutics that target RNA. Starting with BRX857, we showed that a natural phosphate group required for binding could be substituted by a carboxylic group. Subsequent characterization of BRX1555 indicated that a negative charge could be bypassed altogether. Ribocil further demonstrated that an unrelated chemical scaffold supports a similar structural organization of the binding site. In spite of not interacting with A99, WG-3, a more recent discovery at Merck, stabilizes a similar architecture at the binding site, except for A48 at J3/4, which retains a conformation from the unbound state.^50^ Notably, whereas 6–10 hydrogen bonds were observed between the various natural ligands and the RNA, the BRX compound series presented here relies on only 3–4, ribocil makes only one hydrogen bond to the RNA, and no hydrogen bonding is observed for WG-3. Hence, going from natural to synthetic ligands is synonymous with an increase in hydrophobic ligand-RNA contacts. This trend is reminiscent of modified aptamers with increased affinity for protein targets that displayed predominantly hydrophobic contacts at the aptamer-protein interface.^51^ From these observations, we conclude that for specific and productive binding to this riboswitch, drugs need to comply with a set of principles that can be equally fulfilled by various interaction types, emphasizing the importance of hydrophobicity for selectivity (Figure 6). What successful binders to this riboswitch have in common is that they simultaneously stabilize similar conformations for at least five of the six joining regions that interact with the ligand.

**Figure 6.**
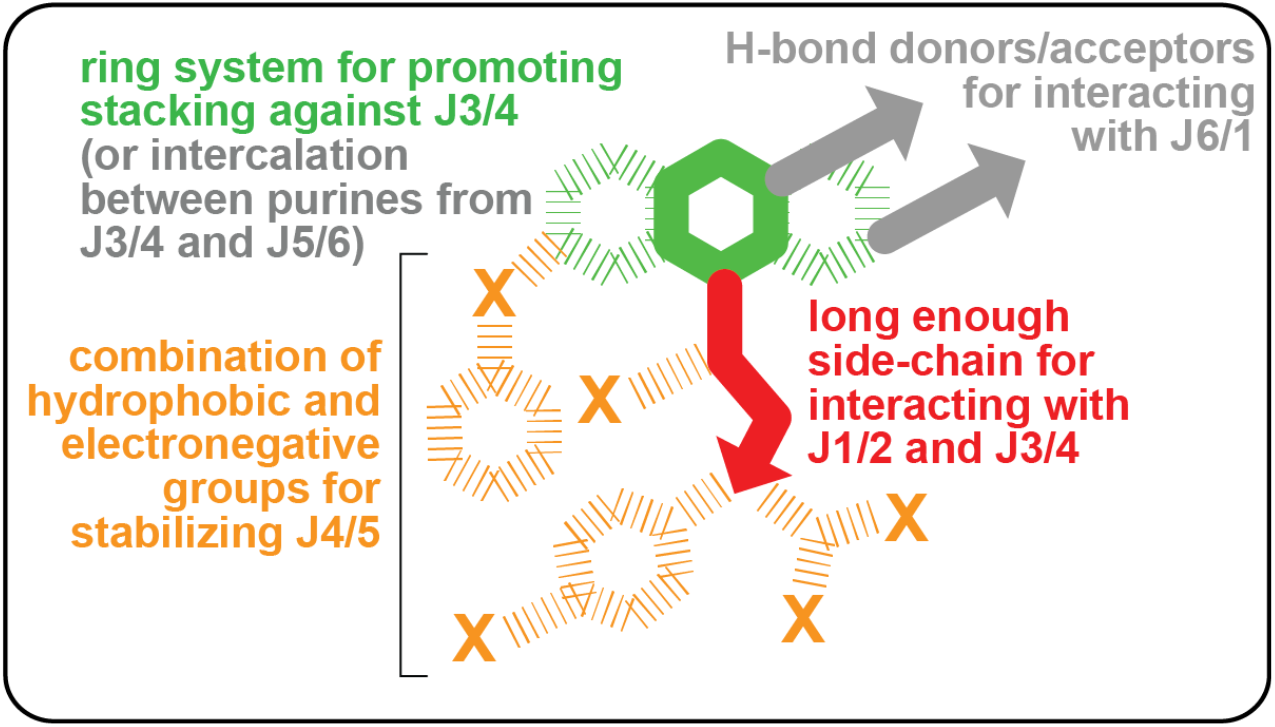
Generic principles for designing drugs that bind to the FMN riboswitch. The most critical aspects of ligand binding are shown in colors around a skeleton schematic of flavin-based chemical structures. Recognition patterns seen with all ligands except WG-3^50^ are shown in gray. Dashed lines indicate putative functionalizations.

Interestingly, a manual search for roseoflavin-like scaffolds within the library that led to the discovery of ribocil produced eight bioactive hits, seven of which contained aryl halides attached to an isoalloxazine heterocycle, some looking very close to 5FDQD.^32^ In particular, the only difference between 5FDQD and the analog named ‘C003’ is the nature of the alkyl chain connecting the isoalloxazine heterocycle to the fluorobenzyl moiety (in C003, the chain contains only one carbon atom). Although C003 and related compounds were shown to have similar limitations to roseoflavin *in vivo*, supplying them to the structure-guided pipeline we describe here would have certainly accelerated the process of discovering 5FDQD. This striking example further attests of the promising future for 5FDQD and its derivatives in drug discovery, while stressing that although structure-based drug design and high throughput screening have both gained enough maturity to advance biomedicine, they could benefit from being further integrated with one another.

## METHODS

### Chemical synthesis

Compounds were either purchased (lumiflavin, 3B Scientific Corporation; roseoflavin, Toronto Research Chemicals; FMN, Sigma-Aldrich) or synthesized (RoFMN and all BRX compounds, see SI Methods). The progress of all reactions was monitored on Merck precoated silica gel plates (with fluorescence indicator UV254) using ethyl acetate/n-hexane as solvent system. Column chromatography was performed with Fluka silica gel 60 (230-400 mesh ASTM) with the solvent mixtures specified in the corresponding experiments. Spots were visualized by irradiation with ultraviolet light (254 nm). Melting points (mp) were taken in open capillaries on a Stuart melting point apparatus SMP11 and were uncorrected. Proton (^1^H) and carbon (^13^C) NMR spectra were recorded on a Bruker Avance 500 (500.13 MHz for ^1^H; 125.76 MHz for ^13^C) using DMSO-d_6_ as solvent. Chemical shifts are given in parts per million (ppm) (δ relative to residual solvent peak for ^1^H and ^13^C). Elemental analysis was performed on a Perkin-Elmer PE 2400 CHN elemental analyzer and a vario MICRO cube elemental analyzer (Elementar Analysensysteme GmbH). IR spectra were recorded on a Varian 800 FT-IR Scimitar series. Optical rotation was given in deg cm^3^ g^-1^ dm^-1^. High-resolution mass spectrometry (HRMS) analysis was performed using a UHR-TOF maXis 4G instrument (Bruker Daltonics). If necessary, purity was determined by high performance liquid chromatography (HPLC) using an Elite LaChrom system [Hitachi L-2130 (pump) and L-2400 (UV-detector)] and a Phenomenex Luna C18 column. Purity of all final compounds was 95% or higher.

### In-line probing assays

The 165 *ribD* RNA used for in-line probing assays was prepared by *in vitro* transcription using a template generated from genomic DNA from *Bacillus subtilis* strain 168 (American Type Culture Collection, Manassas, VA). RNA transcripts were dephosphorylated, 5′ ^32^P-labeled, and subsequently subjected to in-line probing using protocols similar to those described previously.^39^ The *K_D_* for each ligand was derived by quantifying the amount of RNA cleaved in regions where a change was observed (regions 1–6 on Figure S1), over a range of ligand concentrations. The fraction cleaved, x, was calculated by assuming the maximal extent of cleavage is observed in the absence of ligand and the minimal extent of cleavage is observed in the presence of the highest ligand concentration. The apparent *K_D_* was determined by using GraphPad Prism 6 or Kaleidagraph software to fit the plot of the fraction cleaved, *fc*, versus the ligand concentration, [L], to the following equation: *fc* = *K_D_* ([L] + *K_D_*)^-1^.

### *In vitro* RNA transcription termination assays

Single-round RNA transcription termination assays were conducted following protocols adapted from a previously described method.^54^ The DNA template covered the region −382 to +13 (relative to the start of translation) of the *B. subtilis ribD* gene and was generated by PCR from genomic DNA from the 168 strain of *B. subtilis*. To initiate transcription and form stalled complexes, each sample was incubated at 37°C for 10 min and contained 1 pmole DNA template, 0.17 mM ApA dinucleotide (TriLink Biotechnologies, San Diego, CA), 2.5 μM each of ATP, GTP, and UTP, plus 2 μCi 5’-[α-^32^P]-UTP, and 0.4 U *E. coli* RNA polymerase holoenzyme (Epicenter, Madison, WI) in 10 μl of 80 mM Tris-HCl (pH 8.0 at 23 °C), 20 mM NaCl, 5 mM MgCl_2_, 0.1 mM EDTA and 0.01 mg/mL bovine serum albumin (BSA, New England Biolabs). Halted complexes were restarted by the simultaneous addition of a mixture of all four NTPs (at a final concentration of 50 μM) and 0.2 mg/mL heparin to prevent re-initiation, as well as varying concentrations of ligand as indicated to yield a final volume of 12.5 μl in a buffer containing 150 mM Tris-HCl (pH 8.0 at 23 °C), 20 mM NaCl, 5 mM MgCl_2_, 0.1 mM EDTA and 0.01 mg/mL BSA. Reactions were incubated for an additional 20 min at 37 °C, and the products were separated by denaturing 10% polyacrylamide gel electrophoresis (PAGE) followed by quantitation of the fraction terminated at each ligand concentration by using a phosphorimager (Storm 860, GE Healthcare Life Sciences).

### SHAPE probing

Experiments were performed according to published SHAPE protocols.^24, 30, 40^ The *B. subtilis* RNA containing residues 23–166 of the *165 ribD* element^30^ was diluted to a concentration of 2.0 μM in 50 mM HEPES pH 8.0, and incubated for 2 min at 95 °C followed by 3 min at 4 °C. Two parts of this RNA solution were combined to one part of a folding buffer so that the following buffer and ionic concentrations would be reached: 100 mM KCl, 0–15 mM MgCl2 (chosen [MgCl2]: 0, 0.1, 0.5, 2.0, 15.0 mM, 50 mM HEPES pH 8.0, ±10 or 100 μM ligand. This mixture was incubated for 10 min at 37 °C. Two experiments were carried out for each tested compound at each ligand concentration. NMIA modification was performed in the presence of 6.5 mM NMIA for 45 min at 37 °C, at each magnesium concentration specified above. Samples were precipitated for 16 h at −20 °C in 70% EtOH, 100 mM Na-acetate pH 5.3, in the presence of 20 μg of glycogen, centrifuged for 35 min at 13500xg, dried for 10 min under vacuum and resuspended in 9.0 μL 0.5x T.E. buffer (10 mM Tris, 1mM EDTA, pH 8.0). Primer extension by reverse transcription was performed as described,^40^ except that the extension step was performed for 15 min at 50 °C. Primers used for the extension were designed to pair with a binding site in the 3’ structure cassette. Extensions products were resolved by analytical gel electrophoresis, as described.^24^ The gel image (.gel file; Figure S2) was aligned in SAFA v.1.1.^55–56^

### Crystallization

Crystals of complexes with BRX1151, BRX1354 and BRX 1555 were obtained using the hanging-drop vapor diffusion method, following previously published procedures (see SI Methods).^30, 33^ The BRX1354 and BRX1555 complexes were crystallized using RNA sequences and minor variations of the crystallization conditions from the first published structure of the FMN riboswitch^33^ (200 mM MgCl_2_, 100 mM MES-NaOH (pH 6.5), 8.5% w/v polyethylene glycol 4000 (BRX1354); 200 mM MgCl2, 100 mM MES-NaOH (pH 6.5), 8.5% w/v polyethylene glycol 4000, 2.5% DMSO (BRX1555); 20°C), while the BRX1151 complex was crystallized using alternative sequences and crystallization conditions^30^ (320 mM MgCl_2_, 10 mM Tris-HCl (pH 8.4), 8.25% w/v polyethylene glycol 4000; 20 °C). Cryoprotectant was introduced gradually to final concentrations of 10–13.5 % PEG 4000, 10% sucrose and 15% glycerol (BRX1354, BRX1555), or directly to 25% glycerol (BRX1151). Crystals were then harvested with nylon loops and flash cooled by plunging into liquid nitrogen.

### Data collection and processing

Diffraction data for BRX1151 were collected on the X25 beamline at the National Synchrotron Light Source (NSLS) at Brookhaven National Laboratory, as part of the mail-in crystallography program,^57^ and processed via autoPROC^58^ (space group P3_1_21). Diffraction data for BRX1354 and BRX1555 were collected on the X29A beamline at NSLS. Merged data from two (BRX1555) or three (BRX1354) crystals were corrected for anisotropy and rescaled with the Diffraction Anisotropy Server, [http://www.doe-mbi.ucla.edu/~sawaya/anisoscale], indexed in the trigonal space group P3_1_21, and scaled using DENZO and SCALEPACK in the program suite HKL2000.^59^

### Structure determination

For all crystal structures presented, atomic coordinates from the split RNA FMN riboswitch crystal structure PDB code 3F4E were used as a starting model for a molecular replacement search using PHASER.^60^ Ligand, ions and solvent molecules were intentionally omitted from the search model. A solution was successfully obtained in space group P3_1_21 for one riboswitch in an asymmetric unit with high scores (RFZ, TFZ > 10) for the rotation function, translation function, least-likelihood gain and with no atom clashes. A SMILES string for each ligand was generated and supplied to GRADE (Global Phasing, UK) to obtain an atomic model and restraints for refinement in BUSTER-TNT.^61^ After iterative rounds of manual model adjustment and refinement, the ligands were placed in sigma-A weighted F_o_-F_c_ maps. Given the medium resolution of the structures, solvent density was left unassigned, except for density peaks in which we tentatively placed water molecules, anionic, and cationic ions, in agreement with previous rules.^62–65^

## CONTRIBUTIONS

Q.V., K.F.B., P.C. and R.T.B. designed the study. P.A., P.C., J.B., P.W. led the compound design and chemical optimization efforts. J.B., H.K., K.W.K, P.W., and J.W. synthesized and purified RoFMN, BRX247, BRX830, BRX857, BRX897, BRX931, BRX1027, BRX1224 and BRX1354. R.C.G. and H.J.S. synthesized and purified BRX1151 and BRX1555. K.F.B. carried out the functional and in-line probing assays. R.K.S. crystallized the BRX1354 and BRX1555 complexes and solved their structures. Q.V. and E.M. performed the SHAPE mapping experiments, crystallized the BRX1151 complex and solved its structure. Q.V. and F.R. finalized the refinement of all structures. Q.V., K.F.B. and R.T.B. wrote the manuscript. All authors have given approval to the final version of the manuscript.

## ACKNOWLEDGEMENTS

We wish to thank Annie Héroux at Brookhaven National Laboratory for data collection and processing of the BRX1151 complex; Raymond S. Brown, Eric Fontano, and Andre White for contributions to the crystallization and structure determinations of the BRX1354 and BRX1555 complexes; Filip Leonarski and Pascal Auffinger for helpful discussions about solvent; Ron Breaker for support. R.T.B. acknowledges support by a grant from the National Institutes of Health (R01 GM073850). This research used beamline X25 of the National Synchrotron Light Source, a U.S. Department of Energy (DOE) Office of Science User Facility operated for the DOE Office of Science by Brookhaven National Laboratory under Contract No. DE-AC02-98CH10886.

## TABLES

**Table 1.** Data collection and refinement statistics.

|  | BRX1151 | BX1354 | BRX1555 |
| --- | --- | --- | --- |
| **Data collection** |  |  |  |
| Space group | P3<sub>1</sub>21 | P3<sub>1</sub>21 | P3<sub>1</sub>21 |
| Cell dimensions a = b, c (Å); $\alpha = \beta, \gamma$ (°) | 74.4, 140.7; 90, 120 | 71.1, 140.5; 90, 120 | 71.6, 140.3; 90, 120 |
| Wavelength (Å) | 0.979 | 1.5418 | 1.075 |
| Resolution range (Å)<sup>a</sup> | 140.1–3.0 (3.01–3.00) | 20.0–2.8 (2.9–2.8) | 25.0–2.8 (2.9–2.8) |
| Redundancy | 6.8 (6.6) | 7.1 (1.4) | 11.2 (2.7) |
| Completeness (%) | 99.8 (98.9) | 83.4 (50.9) | 96.8 (87.0) |
| $\langle I/\sigma(I) \rangle$ | 17.4 (2.4) | 9.1 (1.7) | 24.6 (2.3) |
| R<sub>merge</sub> (%) | 6.8 (78.1) | 8.5 (38.4) | 7.4 (37.8) |
| **Refinement** |  |  |  |
| Resolution range (Å)<sup>a</sup> | 30.9–3.03 (3.39–3.00) | 19.55–2.88 (3.22–2.88) | 23.10–2.80 (3.13–2.80) |
| No. of reflections | 8941 (2541) | 8313 (1611) | 10388 (2619) |
| R<sub>work</sub> | 0.223 (0.265) | 0.199 (0.304) | 0.193 (0.251) |
| R<sub>free</sub> | 0.242 (0.270) | 0.257 (0.339) | 0.229 (0.282) |
| No. of non-hydrogen atoms; average B-factor (Å<sup>2</sup>) |  |  |  |
| RNA | 2384; 86.8 | 2333; 77.4 | 2337; 82.4 |
| Ligand | 39; 63.6 | 41; 43.4 | 36; 59.6 |
| Solvent | 13 | 15 | 13 |
| RMSD<sup>b</sup> (bonds; Å) | 0.01 | 0.01 | 0.01 |
| RMSD<sup>b</sup> (angles; °) | 0.76 | 0.71 | 0.74 |
| Clashscore (percentile)<sup>c</sup> | 5.2 (100<sup>th</sup>) | 3.1 (100<sup>th</sup>) | 1.7 (100<sup>th</sup>) |
| PDB ID | 6DN1 | 6DN2 | 6DN3 |

<sup>a</sup> Values for the highest resolution shell are given in parentheses.
<sup>b</sup> Root-mean square deviations from ideal values.
<sup>c</sup> Molprobity clashscore is the number of serious steric overlaps ( $> 0.4$ Å) per 1000 atoms (100<sup>th</sup> percentile is the best among structures of comparable resolution).

